# Validation of optical coherence tomography as a tool to identify differentiation key drivers in 3D *in vitro* conjunctival models

**DOI:** 10.64898/2026.01.13.698626

**Authors:** Julian Schwebler, Fabiola Walz, Geraldine Beer, Constantin Berger, Olivier Thouvenin, Djida Ghoubay, Rodrigo Meléndez García, Kate Grieve, Christian Lotz

**Author notes:** Contributed equally to this study (Julian Schwebler, Fabiola Walz). Corresponding author: Christian Lotz **Email:**. **Author Contributions:** J.S., F.W., C.B., and C.L. designed the study. J.S., F.W., G.B., D.G., O.T., and R.M.G. performed the experiments. J.S., F.W., G.B., C.B., D.G., K.G, O.T., and R.M.G. analyzed data. C.L. and D.G. supervised the study and provided material. J.S. and F.W. wrote the manuscript. All authors revised the manuscript and agreed to the submitted version of the manuscript. **Competing Interest Statement:** The authors declare no competing interests.

## Abstract

Conjunctival *in vitro* models present a valuable system to investigate conjunctival tissue homeostasis and pathologies. Combinations of collagen and fibroblasts as a stroma equivalent and the supplementation with serum have been reported to promote the differentiation of epithelial cells. However, how the individual factors affect differentiation of ocular surface cells is insufficiently understood. In this study, we analyzed the effect of serum concentration, a collagen matrix, and fibroblasts on conjunctival differentiation in a 3D *in vitro* model. For this purpose, we developed a computational analysis pipeline for the quantification of optical coherence tomography (OCT) data sets, allowing a time resolved, non-invasive assessment of conjunctiva epithelium differentiation, including goblet cell density. High-resolution dynamic full-field OCT (D-FFOCT) was employed to verify the identity of goblet cells. Conjunctival markers were further analyzed via histology, real-time quantitative PCR, and ELISA to confirm the data of the OCT analysis pipeline. We found that serum is required to induce epithelial differentiation while higher concentrations of 5 – 10% impaired epithelial development. The culture on a collagen matrix increased conjunctival markers upon stimulation with serum, while the co-culture with fibroblasts increased epithelial stratification. Increased serum concentration resulted in the increased occurrence of goblet cells of up to 20 cells/mm². Altogether, the complementary analyses confirmed the quantified OCT data. Summarized, we identified the combination of serum (3%), collagen, and fibroblasts as a condition resulting in the highest physiological resemblance. Altogether, our study emphasizes the need for fine-tuning of culture conditions for 3D *in vitro* models.

**SIGNIFICANCE STATEMENT:** Physiologically relevant in vitro conjunctiva models are essential for studying ocular surface homeostasis and disease. Our study refines 3D conjunctival culture conditions to more closely resemble native tissue by systematically identifying serum concentration, collagen scaffolds and fibroblasts as key drivers of epithelial differentiation. By applying non-invasive optical coherence tomography analysis, we enable longitudinal assessment of tissue maturation, addressing the need for non-destructive, repeatable tissue analysis.

**GRAPHICAL ABSTRACT:** 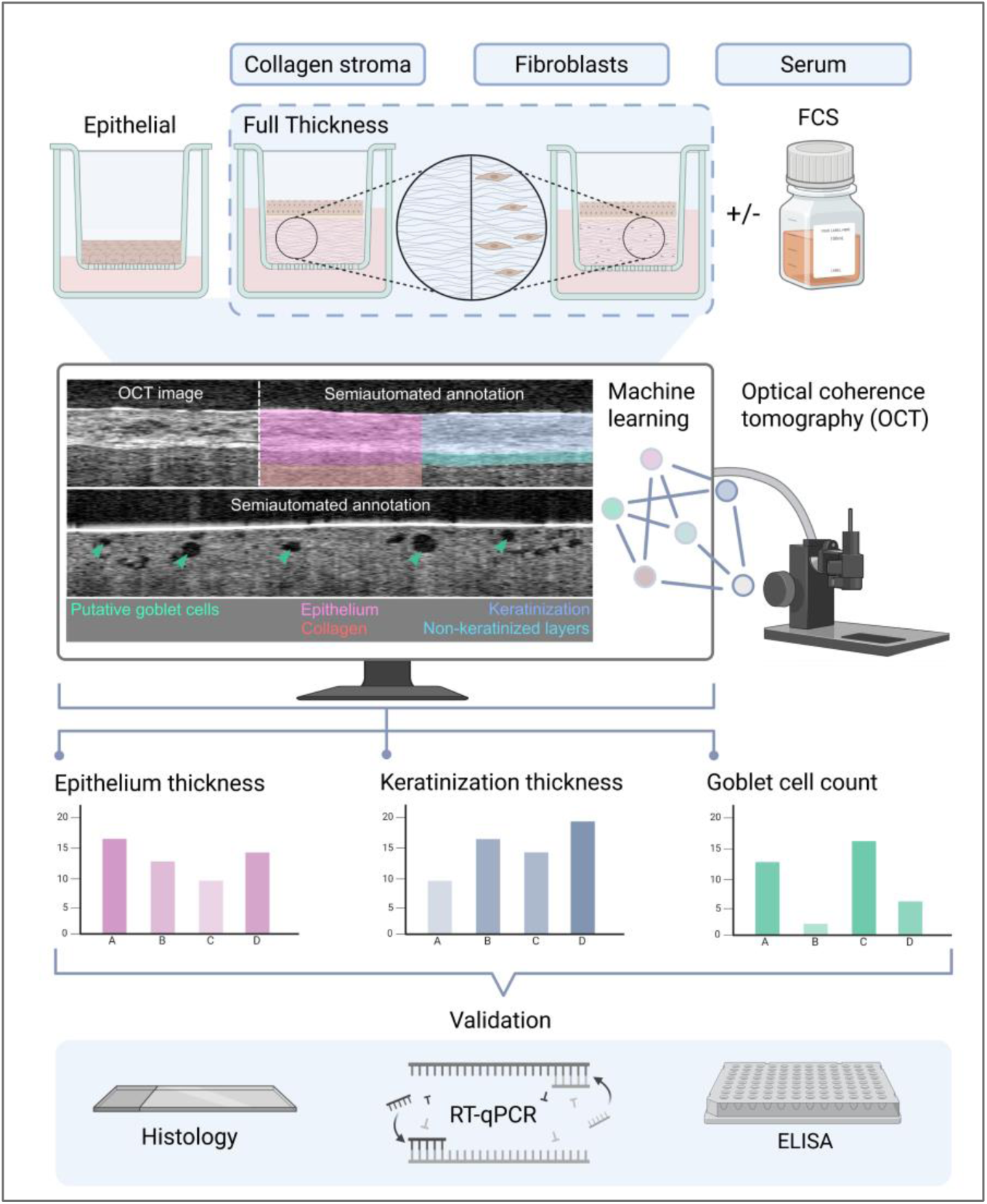

## INTRODUCTION

The conjunctiva plays a crucial role in maintaining the homeostasis of the tear film. Its secretion of mucus stabilizes the tear film by increasing the tear break-up time, while simultaneously protecting the ocular surface from pathogens^1^. The conjunctival epithelium consists of non-keratinized cells that express membrane-associated mucins, as well as specialized goblet cells, which secrete gel-forming and soluble mucins. The most prominent secreted mucin of conjunctival goblet cells is the gel-forming mucin 5AC (MUC5AC)^2–4^.

3D *in vitro* models of conjunctival tissue are used to investigate differentiation mechanisms of epithelial cells into goblet cells as well as to study diseases that affect the conjunctiva such as Dry Eye Disease or infections^5–8^. However, close physiological resemblance is needed for *in vitro* models to increase the transferability of the underlying research questions. In the case of the conjunctival epithelium, this includes a non-keratinized stratified epithelium with differentiated, mucin-expressing goblet cells. One prominent supplement used to generally trigger differentiation processes in *in vitro* models is human or bovine serum, typically at concentrations of 5 – 10%^7–10^. Furthermore, the co-culture of fibroblasts within an extracellular matrix (ECM) scaffold has previously been found to promote the differentiation of conjunctival epithelial cells^8,11^. Studies have shown that fibroblasts modulate epithelial cell behavior and enhance their differentiation^12^, while fibroblast growth factors (FGF) such as FGF2 and FGF10 promote goblet cell differentiation in the conjunctiva and intestine^12–15^. While these studies provided important insights into cell culture and differentiation processes, the specific effects of the individual variables – serum, ECM scaffolds, and fibroblasts – on the differentiation and stratification of epithelial cells is still unresolved.

A factor that is complicating research on *in vitro* models not only of the conjunctiva is the lack of reliable non-invasive imaging techniques. Histological and immunohistochemical staining present the gold standard for visualizing the tissue architecture in 3D *in vitro* models. However, fixation and labelling of the model require the destruction of the model, can introduce technical artifacts such as tissue shrinking, and capture only a snapshot of the model. Optical Coherence Tomography (OCT)^16^ has emerged as a label-free, non-invasive imaging technique overcoming some of the disadvantages of label-dependent imaging techniques. OCT is widely used in ophthalmology to diagnose a variety of pathologies, such as macular degeneration and glaucoma^16,17^. Advancements of OCT have fueled its use in biomedical research such as cancer progression or kidney imaging^18–20^. While offering a limited resolution compared to microscope-based imaging techniques, OCT allows live-recording of *in vitro* models, which reduces the number of required models and allows to investigate spatiotemporal processes. Recently, dynamic full-field optical coherence tomography (D-FFOCT) module coupled to a commercial microscope equipped with a high-numerical aperture objective and a stage-top incubator has enabled imaging of retinal organoids^21^ and other *in vitro* models^22^ at subcellular resolution, revealing 3D structure with high resolution and intrinsic dynamic activity due to metabolic activity and cell function^23–26^.

In this study, we analyzed the effect of various fetal calf serum (FCS) concentrations, a collagen scaffold, and fibroblasts on conjunctival epithelial differentiation. We used OCT imaging to analyze 3D conjunctiva models with respect to goblet cell number, keratinization, and epithelial thickness via OCT. Quantification of these parameters was achieved using the machine-learning-tool of the IMARIS software, providing a new approach of non-invasively evaluating 3D *in vitro* models on a qualitative and quantitative level. We found that all three factors have a distinct effect on conjunctival 3D *in vitro* models. In epithelial models, optimal FCS concentrations induced conjunctival differentiation, including an increased goblet cell number, and led to a disrupted stratification at higher concentrations. The use of a collagen matrix reduced keratinization and increased mucus production, while at the same time, it decreased the FCS concentration required to initially induce epithelial differentiation. Fibroblasts embedded in the collagen matrix led to better stratification of epithelial layers, while no significant goblet cell promoting effect was observed. The findings of this study give new insights into the effects and importance of culture conditions for conjunctival tissue and can be used as orientation to achieve closer physiological resemblance in conjunctival *in vitro* models.

## RESULTS

### Native conjunctiva and tissue models exhibit comparable morphology in OCT analysis

Conventional methods to evaluate and characterize tissue models like histology are usually invasive, resulting in the destruction of the model. OCT has emerged as powerful non-invasive alternative, allowing the continuous monitoring and thus the observation of spatiotemporal processes within a single model^20,24,27^. Within this study, we employed OCT as a non-invasive analysis method for studying the effects of FCS concentrations and the presence of fibroblasts on conjunctival tissue model formation, with a particular focus on epithelial thickness, formation of unphysiological keratinization and goblet cell differentiation. To validate OCT as a suitable method to analyze epithelial conjunctival *in vitro* models, we initially used native human and porcine conjunctival tissue as a benchmark (Fig. 1A). OCT imaging of native tissue allowed the visualization of the conjunctival architecture, including a distinction between the epithelial and stromal compartment. Analysis of the FTConM revealed comparable structures with an observable demarcation of the epithelial layer. Dark, regularly shaped spots within the epithelium were visible in all three samples, which, according to their size and location putatively correlated to mucus-producing goblet cells. AB and PAS staining of the same porcine conjunctival tissue showed mucus filled cells, corroborating that the dark spots in the OCT indeed resemble goblet cells (Fig. 1B). To further validate our assumptions, we performed D-FFOCT on native human conjunctiva and the FTConM (Fig. 1C). In both samples, we detected cells with vacuoles visible as a dark inclusion that closely resemble the morphology of goblet cells.

**Figure 1.**
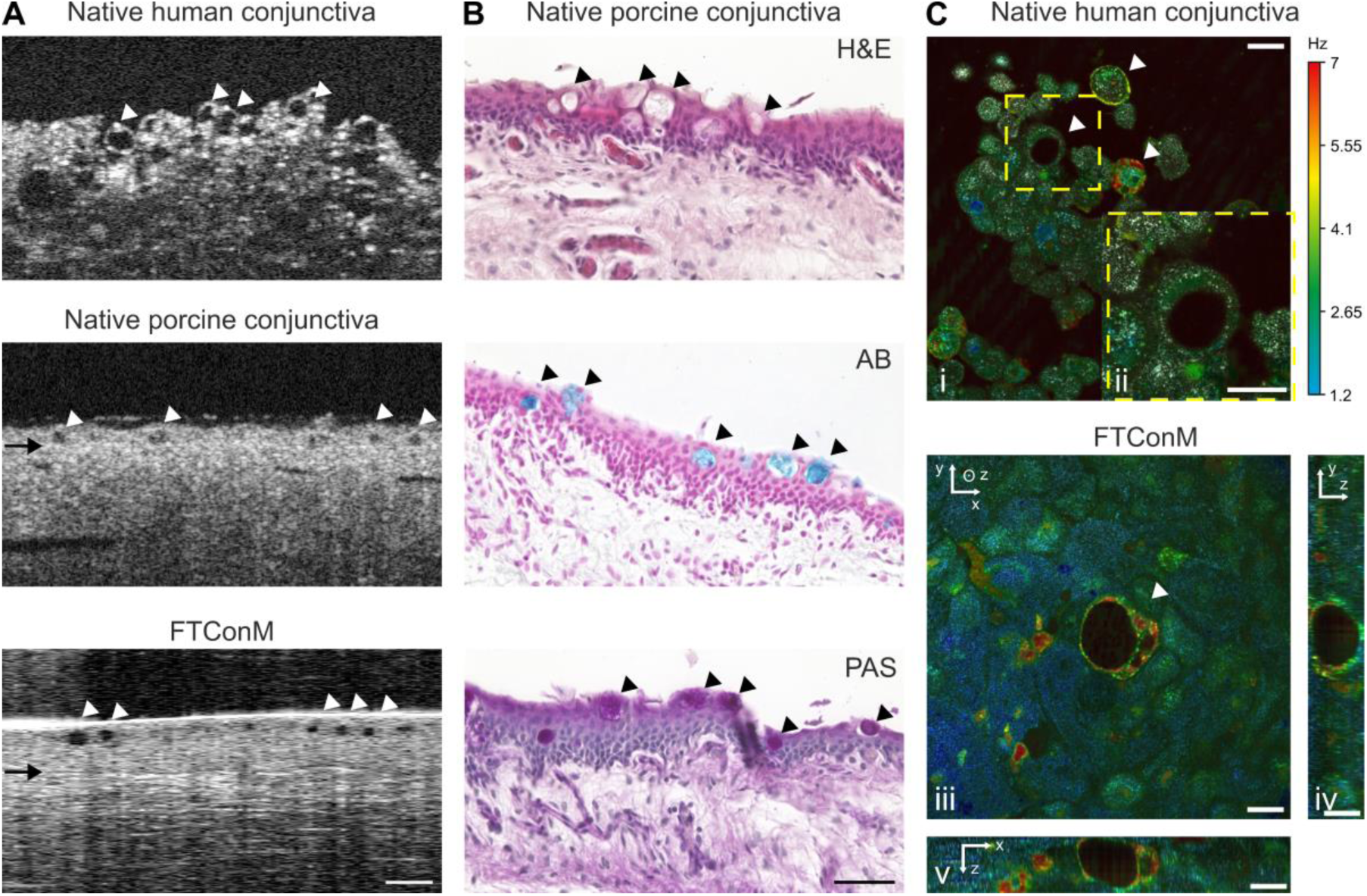
Comparison to *ex vivo* tissue verifies OCT as an adequate method to non-invasively detect epithelial architecture and goblet cells in 3D conjunctival *in vitro* models. **A** OCT imaging of native human and porcine conjunctiva and a FTConM, w/ Fib., 3% FCS. Scale bar:100 µm. Black arrows indicate the transition from epithelium to stroma. **B** Hematoxylin&Eosin (H&E), Alcian blue (AB), and Periodic Acid Schiff (PAS) staining of native porcine conjunctiva. Scale bar: 50 µm. **C** Dynamic full-field OCT (dFFOCT) z-stack of a native human conjunctiva (i, ii) and a FTConM (iii, iv, v) with an axial step of 10 µm for native conjunctiva and 1 µm for FTConM. Hue scales from 1.2 to 7 Hz mean frequency. **i** A selected XY cross-section of native conjunctiva. **ii** zoom-in on (i), indicated by the yellow square, is displayed in (ii), where a goblet cell can be distinguished. **iii** An en face (XY) cross-section from within the FTConM, at 17 µm depth. **iv** A reconstructed YZ cross-section from within the FTConM, using 31 en face images with an axial step of 1 µm. **v** A reconstructed XZ cross-section from within the FTConM, using the same en face images as in (iv). Scale bars:20 µm. Goblet cells are indicated by arrowheads.

Summarized, the data verify OCT as an adequate technique to visualize the conjunctival epithelium architecture including goblet cells, opening up the possibility to non-invasively monitor the differentiation and quality of conjunctival *in vitro* models.

### FCS induces differentiation and influences epithelial thickness in conjunctival epithelial models

Having verified the use of OCT for non-invasively visualizing relevant epithelial parameters, we next aimed to use this system to study the effect of varying FCS concentrations on the differentiation of conjunctival models. For this purpose, reconstructed human Conjunctival Epithelium (rhConE) models were cultured for 15 days and supplemented with increasing FCS concentrations ranging from 1 – 10%.

For the extraction of epithelial parameters from the recorded OCT datasets, we performed semiautomated image segmentation using the machine learning-assisted annotation tool of the Imaris software. This enabled the quantification of total epithelial layer thickness, non-keratinized layers, keratinization and goblet cell differentiation as demonstrated in Fig. 2A.

**Figure 2.**
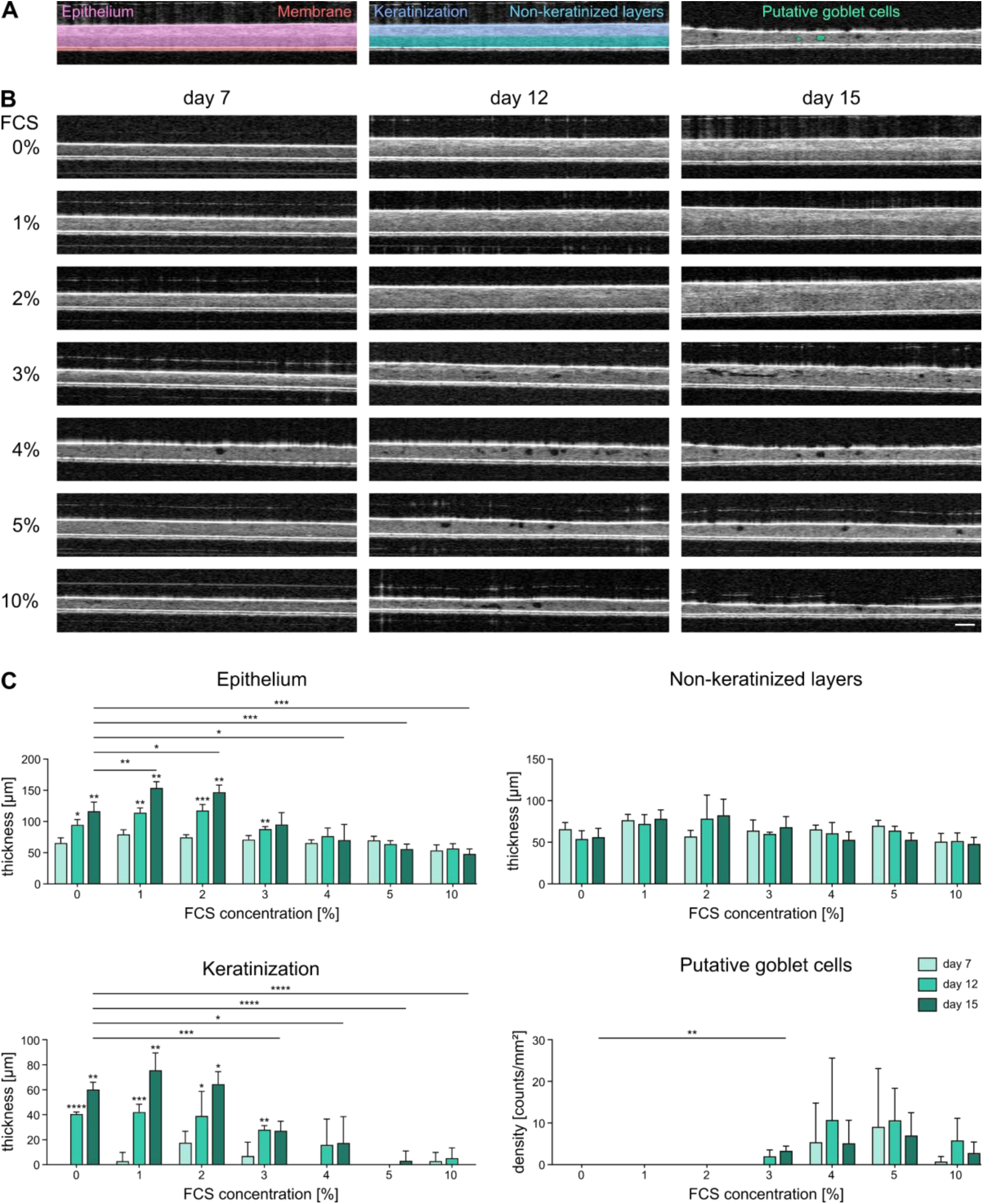
Optical Coherence Tomography (OCT) analysis of rhConE cultured with increasing FCS concentrations. **A** Schematic example of how tissue parameters were analyzed from OCT images. **B** OCT images from day 7, 12 and 15 of rhConE cultured with 0 – 10% FCS. Scale bar: 100 µm. **C** Quantitative analysis of total epithelial thickness, non-keratinized layers, keratinization and goblet cell density of rhConE using OCT 3D stacks and Imaris software (n = 6). Mean + SD; *p < 0.05, **p < 0.01, ***p < 0.001, ****p < 0.0001.

OCT imaging was performed on day 7, 12 and 15 to track the development of the rhConE in presence of various FCS concentrations (Fig. 2B). While no significant differences in total epithelial layer thickness were observed between varying FCS concentrations on day 7 epithelial thickness increased until day 15 in models cultured with FCS concentrations of 0 – 3% (Fig. 2C). Contrarily, higher FCS concentrations led to a slight reduction of epithelial layer thickness. OCT imaging further showed that the changes in epithelial thickness resulted primarily from variations in keratinization, while the thickness of the non-keratinized layers remained unchanged across all conditions and time points. Prominent keratinization was observed for FCS concentrations of 0 – 2% on day 15. Interestingly, further increase of FCS concentrations resulted in a reduction of keratinization, with 3% and 4% FCS showing reduced and 5% and 10% FCS exhibiting strongly diminished keratinization. On day 12, putative goblet cells started to appear in rhConE exposed to 3% FCS or higher, with a maximum number of approximately 10 counts/mm² at 4 – 5% FCS.

To validate the findings made by quantitative OCT imaging, H&E staining was performed on day 15, as well as AB and PAS staining to stain mucus (Fig. 3). RhConE showed a multi-layered epithelium with a thick keratinization at 0 – 2% FCS supplementation. While PAS staining intensified from 0 – 2%, no goblet cells were observed. In correlation with the occurrence of dark spots in the OCT, AB and PAS-positive goblet cells with a cup-like structure were detectable at 3 – 10% FCS. Histological stainings confirmed the reduction of keratinization and epithelial thickness measured in OCT-analyses with increasing FCS concentration. At FCS concentrations of ≥ 4%, we observed a decreased tissue stratification with flattened basal cells, indicating a negative influence of high FCS concentrations on the epithelial organization of rhConE.

**Figure 3.**
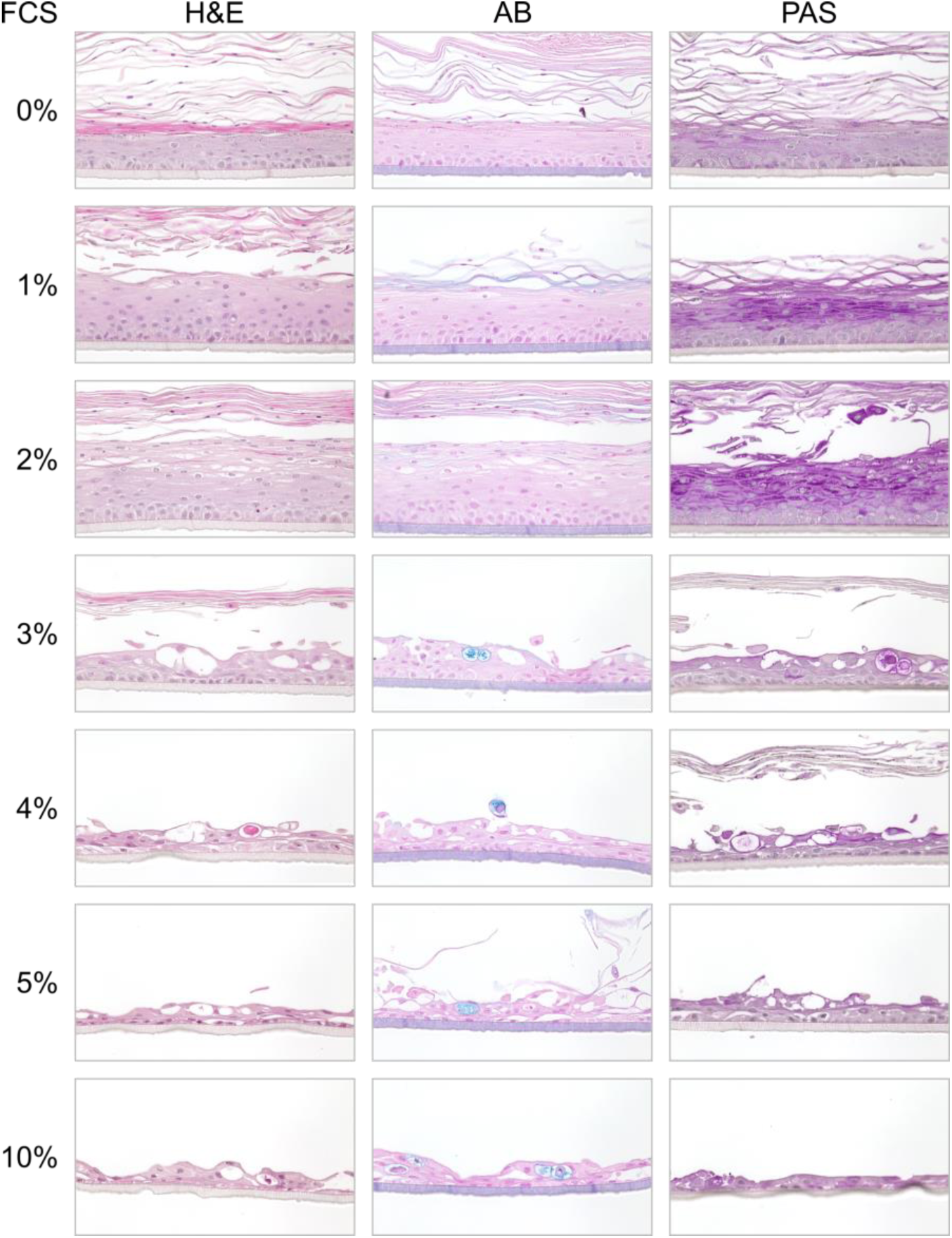
Dose-dependent effects of FCS on the epithelial stratification and presence of mucus-containing cells in rhConE. Hematoxylin&Eosin (H&E), Alcian blue (AB), and Periodic Acid Schiff (PAS) staining of rhConE supplemented with 0 – 10% FCS. Models were stained at day 15 of culture. Scale bar: 50 µm.

Next, we analyzed the gene expression of the conjunctival markers to study the dose-dependent effect of FCS on conjunctival differentiation*. KRT13* expression was significantly elevated at 1% and 2% FCS compared to 0% FCS, with fold changes of 240 ± 105 and 506 ± 72, respectively (Fig. 4A). At ≥ 3% FCS, *KRT13* expression was elevated up to 79-fold compared to rhConE cultured without FCS, but significantly less increased compared to 2% FCS. *KRT19* expression increased in a dose-dependent manner from 0 – 3% and remained constant at ≥ 3% FCS at approximately 1200- to 1500-fold change compared to 0% FCS. In contrast to *KRT19* expression, *LORICRIN*, a marker for the cornified envelope^28^, decreased with increasing FCS concentrations and showed a significant reduction at ≥ 3% compared to the FCS-free condition, indicating a diminishing effect of elevating FCS concentrations on keratinization and supporting the findings made by the OCT data quantification.

**Figure 4.**
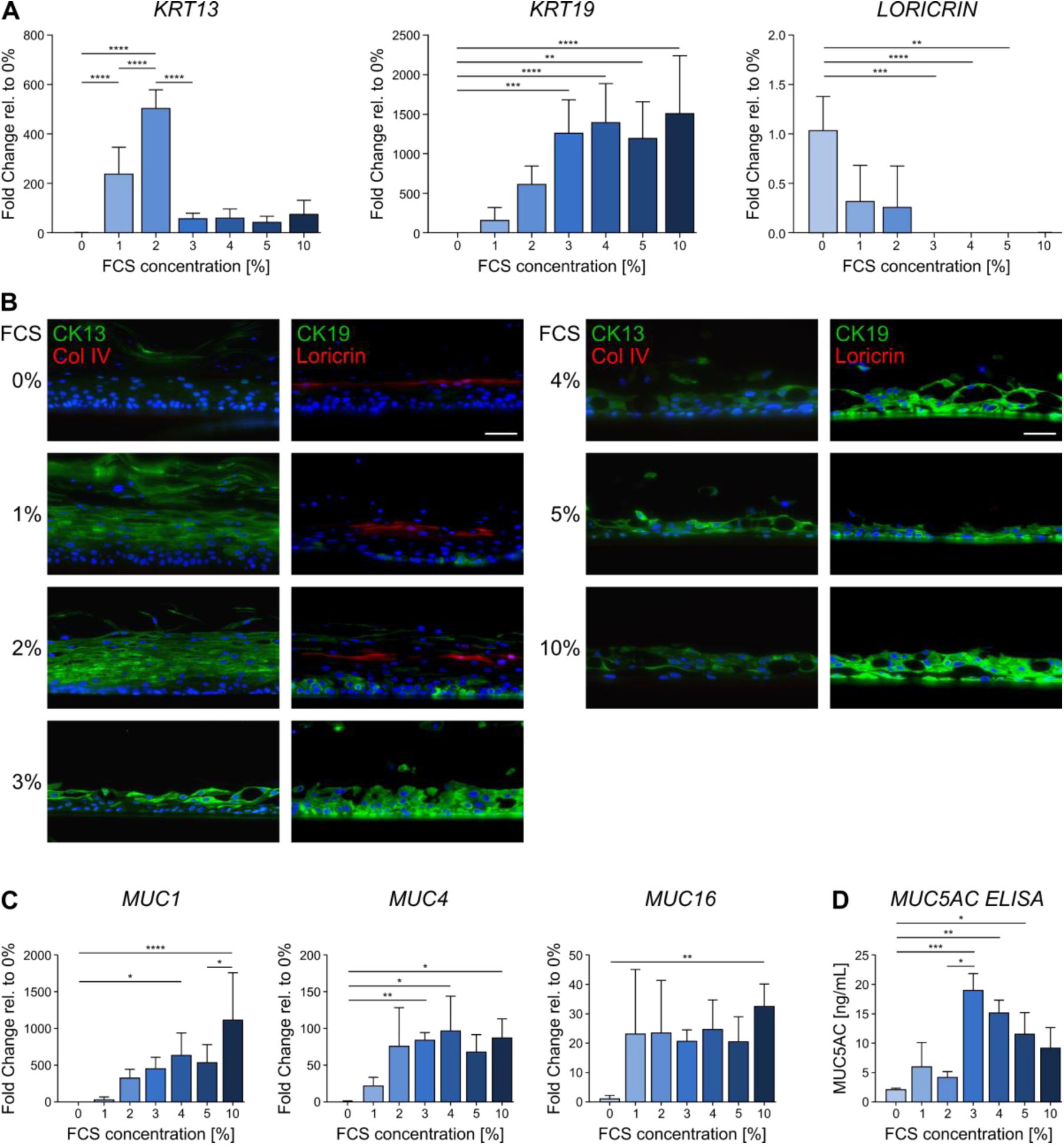
Analysis of specific marker expression in rhConE cultured with FCS concentrations of 0 – 10%. **A** RT-qPCR of KRT13, KRT19, and LORICRIN (n = 3). **B** Immunofluorescence of CK13, CK19, Loricrin, and Col IV. DAPI = blue. Scale bar: 50 µm. **C** RT-qPCR of membrane-associated mucins MUC1, MUC4, and MUC16 (n = 3). **D** ELISA of secreted MUC5AC (n = 3). Mean + SD; *p < 0.05, **p < 0.01, ***p < 0.001, ****p < 0.0001.

Immunofluorescence staining confirmed the gene expression data (Fig. 4B). While the signal of CK13 peaked at 3% FCS supplementation, CK19 staining appeared stronger with increasing FCS concentration. Loricrin staining was detectable in the upper region of the epithelial layers in rhConE supplemented with 0 – 2% FCS, but not detectable at 3 – 10% FCS supplementation, proving a keratinization-inhibiting effect of increasing FCS concentrations. Collagen IV (Col IV), stained for the comparison with FTConM, was not detectable in rhConE.

To verify the beneficial effect of increasing FCS concentrations on the number of goblet cells detected by OCT and histological stainings, gene expression of the membrane-associated mucins *MUC1*, *MUC4*, and *MUC16* was analyzed (Fig. 4C). The expression of all three mucins increased with higher FCS concentration. A significant elevation of *MUC4* expression was measured in models cultured with 3% FCS, while *MUC1* and *MUC16* transcript levels were significantly increased at 4% FCS and 10% FCS, respectively. Furthermore, the secretion of MUC5AC was measured via ELISA (Fig. 4D). The highest MUC5AC secretion was observed at 3% FCS and showed lower concentrations with increasing FCS supplementation, indicating that higher FCS concentrations were detrimental for MUC5AC secretion of goblet cells.

Summarized, the acquired data demonstrate that increasing FCS concentrations prevent keratinization and increase the number of goblet cells of rhConE, while exposure to ≥ 4% FCS impairs a regular epithelial stratification. Importantly, the conducted experiments verified quantitative OCT as an adequate method to assess epithelial architecture and goblet cell number.

### The connective tissue equivalent promotes differentiation and stratification in conjunctival *in vitro* models

As a next step, we analyzed whether the addition of a connective tissue equivalent promotes epithelial cell differentiation and improves the structure of conjunctival models. Conjunctival epithelial cells were cultured on a compressed collagen matrix without (w/o Fib.) or with (w/ Fib.) embedded conjunctival fibroblasts. To investigate the effect of FCS on epithelial formation in the presence of fibroblasts, both conditions were subjected to 0 – 3% FCS supplementation.

OCT-images of FTConM were analyzed using machine learning-assisted segmentation, as performed for rhConE (Fig. 5A). Total epithelial thickness, non-keratinized layers, keratinization and goblet cells could be identified as distinguishable structures (Fig. 5B). Quantitative analyses revealed an increasing epithelial thickness from day 7 (79 ± 7 µm) to day 15(155 ± 19 µm), regardless of FCS concentration (Fig. 5C). In contrast to the rhConE models, keratinization of the epithelium was prevented at lower FCS concentrations, and the total epithelial thickness was not reduced at higher FCS concentrations. While at 0 – 1% FCS the increase of the total epithelial thickness resulted almost exclusively from a thickening of the keratinized layer, FTConM exposed to 2 – 3% FCS remained non-keratinized and exhibited an increased height of the non-keratinized epithelial layers. Interestingly, models cultured with fibroblasts showed fewer epithelial layers at d15 compared to models cultured without. On the other hand, models cultured with fibroblasts showed a more defined stratification especially at higher FCS concentrations, with a clear demarcation between epithelium and collagen. Goblet cells started to appear in models cultured with 2% FCS and, more frequently, with 3% FCS. In both cases, goblet cell frequency increased over time.

**Figure 5.**
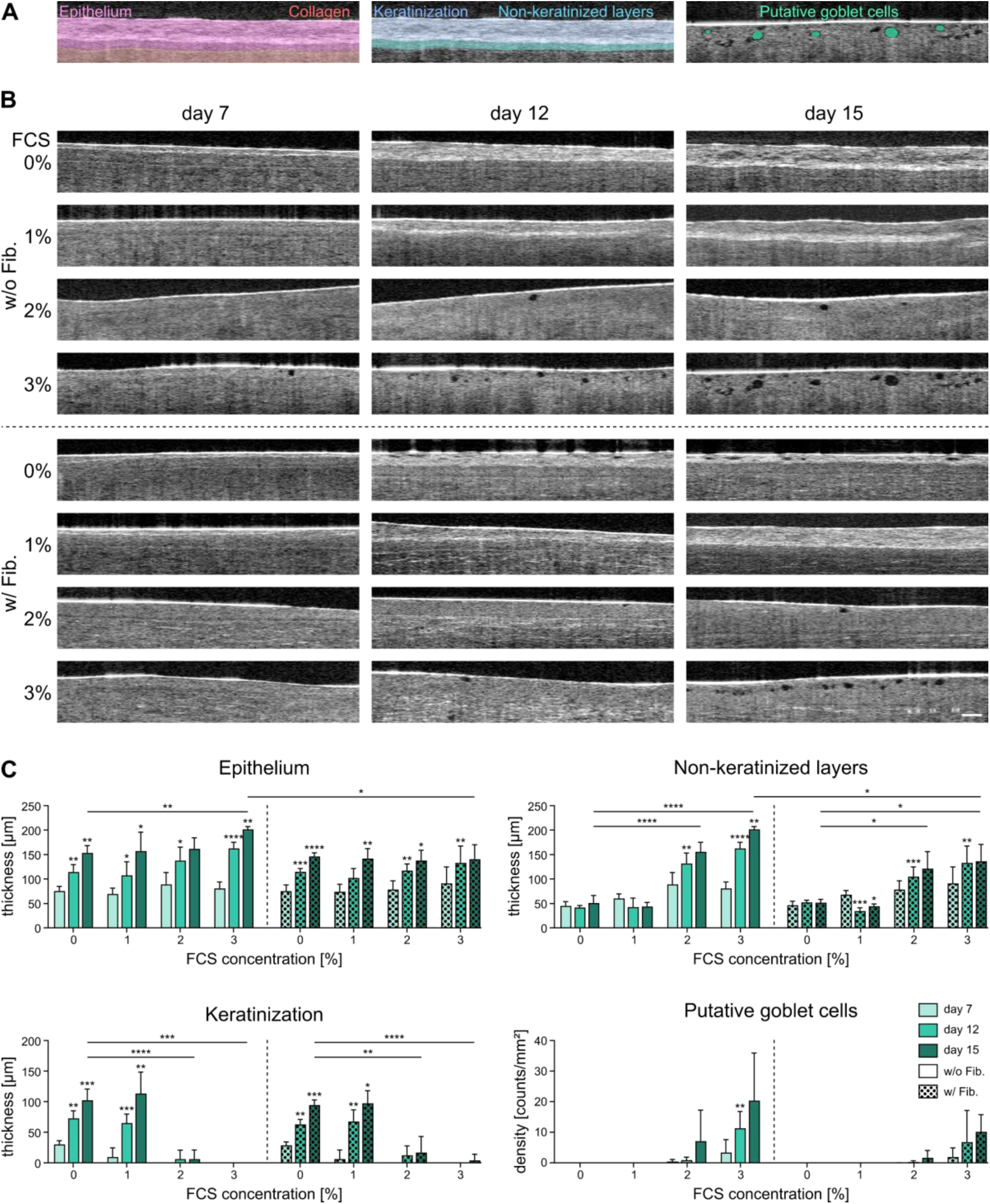
OCT analysis of FTConM cultured without (w/o Fib.) and with (w/ Fib.) fibroblasts and increasing FCS concentrations. **A** Schematic example of how tissue parameters were analyzed from OCT images. **B** OCT images from day 7, 12 and 15 of FTConM cultured with 0 – 3% FCS. Scale bar:100 µm. **C** Quantitative analysis of total epithelial thickness, non-keratinized layers, keratinization and goblet cell density of FTConM using Imaris software (n = 6). Mean + SD; *p < 0.05, **p < 0.01, ***p < 0.001, ****p < 0.0001.

Histological analysis confirmed a reduced keratinization of models with and without fibroblasts cultured at ≥ 2% FCS (Fig. 6). PAS staining gradually intensified from 0% to 3% FCS. Goblet cells were present in FTConM treated with 2% and 3% FCS. In FTConM cultured without fibroblasts, we observed an invasive behavior of epithelial cells into the collagen scaffold. This effect was more prominent with increasing FCS concentrations compared to 0% FCS, where no invasion was detected at all. In models cultured with conjunctival fibroblasts, no invasive epithelial cells were observed. FTConM exposed to 3% FCS exhibited a more distinct basal layer of epithelial cells when cultured with fibroblasts than without. Overall, the stratification of epithelial layers was more structured in models cultured with fibroblasts and higher FCS concentrations, while this was not the case for models without FCS supplementation, suggesting that conjunctival fibroblasts secrete factors that promote epithelial stratification.

**Figure 6.**
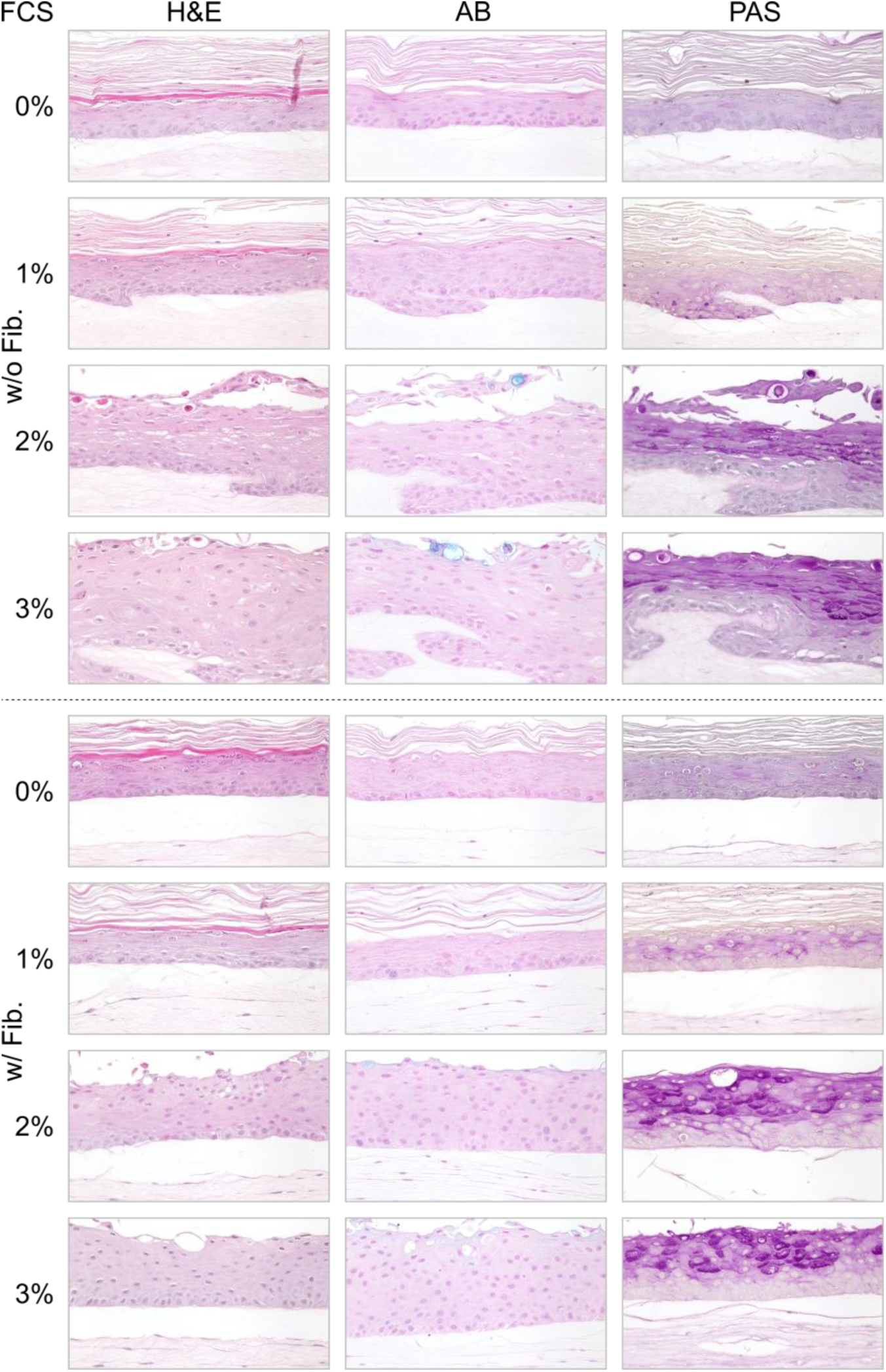
Effects of varying FCS concentrations in FTConM with and without fibroblasts on epithelial stratification and the occurrence of mucus-containing cells. Hematoxylin&Eosin (H&E), Alcian blue (AB), and Periodic Acid Schiff (PAS) staining of FTConM without (w/o Fib.) and with (w/ Fib.) fibroblasts, supplemented with 0 – 3% FCS. Models were stained on day 15 of culture. Scale bar: 50 µm.

Gene expression analysis revealed a significant increase of *KRT13* expression at 2% and 3% FCS, and *KRT19* at 3% FCS, both with and without fibroblasts (Fig 7A). *LORICRIN* was significantly reduced at 3% FCS for both conditions. At 2% FCS, the expression of *KRT13* was significantly higher in models cultured without fibroblasts which was not the case for other FCS concentrations. No significant difference in the expression of *KRT19* and *LORICRIN* was observed between models cultured with and without fibroblasts.

**Figure 7.**
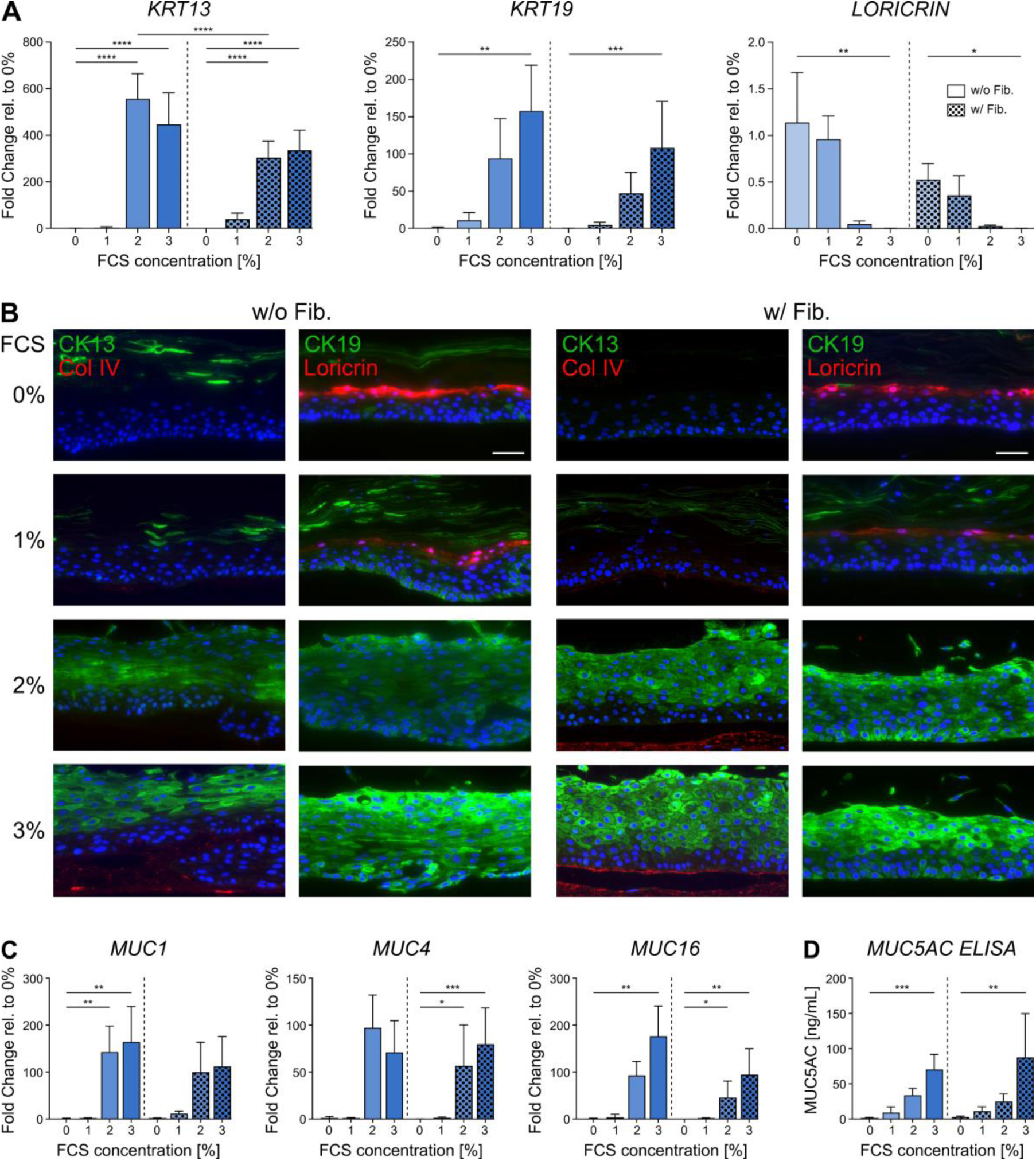
Analysis of specific marker expression in FTConM cultured with FCS concentrations of 0 – 3%. **A** RT-qPCR of KRT13, KRT19, and LORICRIN (n = 3). **B** Immunofluorescence of Cytokeratin (CK) 13, CK19, Loricrin, and Col IV. DAPI = blue. Scale bar: 50 µm. **C** RT-qPCR of membrane-associated mucins MUC1, MUC4, and MUC16 (n = 3). **D** ELISA of secreted MUC5AC (n = 3). Mean + SD; *p < 0.05, **p < 0.01, ***p < 0.001, ****p < 0.0001.

Immunofluorescent staining of FTConM with and without fibroblasts cultured without FCS showed no CK13 and CK19 staining, whereas Loricrin was detectable in both conditions (Fig. 7B). CK13 and CK19 were detectable in FTConM treated with 2% FCS, with an increased signal intensity at 3% FCS, both with and without fibroblasts. The combination of fibroblasts and FCS further resulted in the secretion of the basement membrane protein Col IV. In FTConM without fibroblasts, Col IV was absent at 0 – 2% FCS while single spots were stained at 3%. In FTConM cultured with fibroblasts, small spots of Col IV were detected underneath the epithelial basal layer at 1% FCS, with an increased signal intensity at 2%. With 3% FCS, a defined basement layer of Col IV could be detected underneath the epithelium. In line with the increased number of goblet cells revealed by quantitative OCT, the number of transcripts encoding for the membrane associated mucins *MUC1*, *MUC4*, and *MUC16* was increased by FCS in a dose-dependent manner (Fig. 7C). Similarly, FCS administration resulted in an elevated secretion of the secretory mucin MUC5AC (Fig. 7D). The inclusion of fibroblasts had no effect on mucin expression or secretion.

Taken together, the data confirm that FCS concentrations of 2% and 3% resulted in a more differentiated, non-keratinized epithelium with a thicker epithelial layer and a higher frequency of goblet cells, while the combination with fibroblasts improved epithelial stratification and Col IV deposition.

## DISCUSSION

Previously, we demonstrated that the incorporation of a connective tissue equivalent promotes the differentiation of our conjunctival *in vitro* model towards a physiological morphology^11^. In this study, we analyzed the individual components of the connective tissue equivalent regarding their effects on conjunctival differentiation. We found distinct effects for FCS, collagen, and fibroblasts on epithelial differentiation and stratification. Previous studies demonstrated the promoting properties of fibroblasts and collagen on mucosal differentiation^8,29^. While these studies gave important insights on conjunctival differentiation, FCS was used in their culture medium, making it difficult to determine the effect of individual factors. In general, serum is an important supplement for the culture of various cell types and is widely used at concentrations ranging from 5 – 10%^7–10^. In our study, we show that these concentrations are not ideal for the culture of conjunctival 3D *in vitro* models, as serum concentrations of ≥ 5% prevented the growth of a multi-layered and stratified epithelium. However, in both rhConE and FTConM, FCS was needed to initially induce mucous differentiation and shift conjunctival epithelial cells from a pathological keratinized state toward a more natural conjunctival phenotype. Of the numerous factors contained in FCS, retinoic acid (RA)^30^ and Interleukin-13 (IL-13)^31^ present interesting candidates for mediating the observed effects. In conjunctival organoids, RA has recently been shown to promote differentiation while IL-13 is known to enhance goblet cells and mucus expression in mucosal tissue^32–35^.

The FCS-driven differentiation could also explain the impairment of epithelial stratification observed at higher FCS concentrations. Since cell proliferation decreases with increased differentiation^36^, factors in FCS that cause differentiation may in turn reduce cell proliferation, impairing stratification due to a reduced cell mass. In the light of these findings, fine tuning of FCS or the addition of a defined growth factor supplementation emerges as a vital task to ensure the formation of conjunctival *in vitro* models that properly mimic an *in vivo* histology.

The comparison of rhConE and FTConM showed that besides FCS, collagen had a positive effect on epithelial cells. In detail, we demonstrate in our study that the use of a collagen matrix did not induce differentiation alone but reduced the FCS concentration needed to achieve a conjunctival phenotype, determined by the reduction of keratinization, retainment of a multi-layered epithelium, and induction of goblet cells at lower concentrations. In this way the use of collagen also circumvents the negative effects caused by high serum concentrations. Zhou et al. showed that Col I promoted the expression of MUC5AC^37^ which we could confirm in our study via the comparison of rhConE and FTConM in a MUC5AC ELISA. These beneficial effects of collagen might originate from its lower stiffness compared to cell culture inserts, as well as the interaction between collagen proteins and integrins on epithelial cells, which can induce intracellular downstream signaling^38,39^.

A clear demarcation between the stroma equivalent and the basal layer was lost with increasing FCS concentrations and epithelial cells showed an invasive behavior. Serum, particularly the contained factors TGF-β and EGF, have been shown to promote epithelial-mesenchymal transition (EMT) in epithelial cells^40,41^, characterized by a transition towards more invasive behavior, cell migration and a reduction of cell-cell adhesion markers like E-cadherin^40^. Epithelial invasion was mitigated by the inclusion of fibroblasts into the collagen scaffold. The addition of fibroblasts further resulted in a clear demarcation at all FCS concentrations and an increased stratification of the epithelial layers. The interaction of epithelial and stromal cells is an important factor, especially due to the expression of growth factors and matrix remodeling. Mesenchymal cells express FGFs, of which FGF2, FGF7, and FGF10 promote proliferation^42–45^. However, to our knowledge little is known about the spectrum of secreted growth factors by conjunctival fibroblasts. Thus, future investigations could elucidate which specific interactions contributed to the retainment of epithelial architecture. Furthermore, fibroblasts are known to change and remodel their ECM environment, for example by the secretion of matrix metalloproteinases (MMPs)^46–49^, leading to a contraction of collagen which results in a denser, more structured matrix^50^. By remodeling the collagen scaffold, fibroblasts therefore possibly prevented epithelial invasion in a mechanical manner. We also detected Col IV, an essential component of basement membranes^51,52^, at the interface of scaffold and epithelium which was most defined in the presence of fibroblasts at 2% and 3% FCS supplementation. This might result from the activation and enhanced collagen deposition of fibroblast induced by FCS^53^. As a functional basement membrane is important for the regulation of cell migration and tissue organization^51^, the presence of a developing basement membrane at the scaffold-epithelium interface may have contributed to the improved epithelial stratification observed in our model. Regarding mucus expression, the effect of fibroblasts is unclear in literature. Tsai et al. demonstrated that conjunctival fibroblasts promote mucus producing cells^8^ while Massaro et al. showed in an intestinal co-culture model of fibroblasts and epithelial cells a fibroblast-dependent reduction of gel-forming mucin expression^54^. Interestingly, further studies show that Col IV reduces MUC5AC in airway cells^55,56^. As we observed no effect of fibroblasts on mucus production, further investigation is necessary in the future to determine when fibroblasts influence its expression and whether the presence of Col IV has a respective effect in 3D *in vitro* models.

Within this study, we used OCT to monitor the tissue development over time and to analyze the aforementioned cellular and structural effects. We were able to distinguish model structures, including keratinized and non-keratinized epithelia, as well as the membrane or the stroma equivalent. Effects of culture conditions observed in OCT were supported by RT-qPCR, histological analyses, and ELISA, confirming the validity of this approach. OCT is widely applied for ophthalmologic diagnostics, such as for retinal and ocular surface imaging^57^. Recently, OCT is increasingly used in tissue engineering and *in vitro* applications^24,58,59^. Conventional OCT enabled quantification of vacuoles, which with a very high level of confidence correspond to individual goblet cells. A study by Aguirre et al. confirms the observation of goblet cells as distinct, non-scattering inclusions in optical coherence microscopy images^60^. Although intensity-based image segmentation allowed reliable detection of dark spots within the epithelium, unambiguous identification of goblet cells was complicated by the presence of gaps in the epithelium, which likely result from reduced cell-cell-adhesion and similarly present as dark areas. Inclusion of defined post-segmentation object filtering nevertheless enabled a representative assessment of goblet cell numbers and the differentiation status, supported by the strong correlation between OCT-based quantification and the data from RT-qPCR, ELISA as well as histological analyses. Future possible improvements to maximize the read-out quality of goblet cells include combinatorial approaches, such as additional D-FFOCT images to identify cell structures around vacuoles, as well as dynamic microscopic OCT, which has previously been shown to allow the identification of individual cells^61^. Furthermore, Kim et al. showed the use of moxifloxacin to label goblet cells non-invasively, which could also be used as a complementary method^62^.

Nevertheless, we show that OCT is a suitable method for non-invasive monitoring of conjunctival *in vitro* tissue, as it captures relevant structural features and does not rely on the administration of any substances, enabling a quick evaluation of generated models with high quality. A future step in goblet cell analysis could be the distinction between mucus filled and empty cells. This could enable easier investigation of conjunctival pathologies that affect mucin secretion such as dry eye disease or allergic conjunctivitis^61,63,64^.

Summarized, we used qualitative and quantitative OCT techniques in combination with gene and protein expression analyses to show the individual effects of FCS concentrations, a collagen matrix, and co-cultured fibroblasts on conjunctival differentiation and stratification. Overall, the highest physiological resemblance was achieved by the combination of all three factors with 3% FCS supplementation. This could pave the way for more predictive and relevant *in vitro* models as well as for new non-invasive analysis tools to research differentiation processes.

## MATERIALS AND METHODS

### Conjunctival tissue

The experiments performed in this study followed the Declaration of Helsinki. The local ethics committees (IRB of the Medical Faculty of the University of Wuerzburg; approval numbers 187/17, 195/13, 82/10 and 2018-280_5-dvh and French Society of Ophthalmology, approval number IRB 00008855) approved the use of human material. Conjunctival biopsies were obtained from adult patients of the Department of Ophthalmology, University Hospital Würzburg with informed consent or from enucleated eyes for corneal graft preparation from deceased organ donors, after informed consent from their relatives. The biopsies used for this study did not show other ocular pathologies that affect the conjunctiva such as pemphigoid, pterygium, or significant inflammation. All biopsies were anonymized and assigned to a number.

Porcine tissue was kindly provided by a local slaughterhouse and used immediately upon obtainment.

### Isolation and culture of primary human conjunctival cells

The isolation of primary human conjunctival epithelial cells and fibroblasts as well as the generation of reconstructed human Conjunctival Epithelium models (rhConE) and Full Thickness Conjunctiva Models (FTConM) was performed as previously described^11^. RhConE were cultured in Pro10^-^ Medium (PromoCell Keratinocyte Growth Medium 2 + Supplement Mix (PromoCell, Heidelberg, Germany) + 1% Penicillin/Streptomycin (Sigma Aldrich, Darmstadt, Germany) + 1.5 mM CaCl2 (Sigma Aldrich, Darmstadt, Germany) + 73 µg/mL Ascorbic acid-2-phosphate (Sigma Aldrich, Darmstadt, Germany) + 10 ng/mL Keratinocyte Growth Factor (KGF, PeproTech Proteins, Thermo Fisher, Waltham, USA) + 10% FibroLife medium (without supplement LifeFactor, Epidermal Growth Factor, Transforming Growth Factor-b, Gentamicin, and Amphotericin, Lifeline Cell Technology, San Diego, USA)). Depending on the experiment, FCS was added at concentrations of 1 – 5% or 10%. Epithelial cells were used in passage (p) 3 for model generation while fibroblasts were used in p5.

### Quantitative Optical Coherence Tomography

Optical coherence tomography was used as a non-invasive method to observe and quantify the structural development of the *in vitro* models. To acquire OCT image data, a commercial spectral-domain OCT (SD-OCT) Ganymede system (Thorlabs GmbH, Bergkirchen, Germany), equipped with a GAN621 base unit and an OCT-LK3-BB objective, was used. The light source has a nominal center wavelength of 900 nm and an axial resolution of 3 µm.

On day 7, 12 and 15, volumetric OCT-scans were acquired for each model, with an X-Y scan area of 5 x 2 mm^2^, a pixel size of 10 x 10 µm^2^ and a 25 kHz A-Scan rate.

To analyze the volumetric image data in terms of quantification of epithelial thickness, keratinization and number of goblet cells, the respective structures were annotated using machine learning segmentation in Imaris (Version 10.2.0, Bitplane AG, Zurich, Switzerland). For quantification of the epithelial layers and keratinization, the volume of the annotated structure was divided by the scan area to obtain the average thickness of each layer per scan. The number of goblet cells was approximated by counting dark spots within the epithelium, with a diameter of 20 – 60 µm and a sphericity greater than 0.8.

### Dynamic Full-Field Optical Coherence Tomography

Native human conjunctiva and FTConM were obtained or generated as previously described^11^. Samples were mounted in black, flat-bottom glass 24-well plates compatible with cell culture and microscopy (Cellvis, P24-1.5H-N) and placed in a stage-top incubator (H201-K-FRAME, H201-MW-HOLDER, and OBJ-COLLAR-2532; Okolab). Prior to imaging, samples were equilibrated in the incubator to stabilize temperature and CO₂ concentration (37 °C, 5% CO₂). Dynamic, non-invasive, label-free live-cell imaging was performed using a custom D-FFOCT module coupled to a commercial microscope, as previously described^22^. Volumetric acquisitions were performed with sufficient lateral and axial coverage to resolve goblet cell morphology. Image and volume reconstruction was carried out using ImageJ software.

### Histology

Native tissue as well as conjunctival models were fixed in Rotifix® (Carl Roth, Germany) for 2 h at room temperature (RT) and embedded in paraffin. Sections of 3.5 µm were prepared from paraffin blocks and used for staining. For histological staining, sections were dewaxed and rehydrated before staining with Hematoxylin&Eosin (H&E; Morphisto, Offenbach am Main, Germany), Alcian blue (AB; Morphisto, Offenbach am Main, Germany), and Periodic acid Schiff (PAS; Carl Roth GmbH + Co. KG, Karlsruhe, Germany) according to the manufacturer’s guidelines. Stained sections were dehydrated and mounted with Entellan (Merck, Darmstadt, Germany).

For immunofluorescence, paraffin sections were deparaffinized and rehydrated. Next, heat mediated antigen retrieval was performed by incubation in citrate buffer pH 6 at 95 °C for 20 min. Subsequently, sections were blocked in 5% Bovine Serum Albumin (BSA, Carl Roth, Karlsruhe, Germany) for 20 min. Primary antibodies were applied over night at 4 °C. On the next day, sections were washed three times in washing buffer (PBS^-^ + 0.5% Tween-20, VWR, Radnor, PA, USA). Secondary antibodies were applied for 1 h at RT. Subsequently, the slides were washed three times in washing buffer and mounted with FluoroMount^TM^ containing DAPI (Invitrogen, Carlsbad, CA, USA). The following primary antibodies were used: Cytokeratin 13 (Dilution 1:200, sc-390982, Santa Cruz Biotechnology, Dallas, TX, USA), Cytokeratin 19 (Dilution 1:200, ab7754, Abcam, Cambridge, UK), Collagen IV (Dilution 1:500, ab6586, Abcam, Cambridge, UK), and Loricrin (Dilution 1:500, ab85679, Abcam, Cambridge, UK). The following secondary antibodies were used: AlexaFluor® 555, 647 (dilution 1:400, LifeTechnologies, Carlsbad, CA, USA).

### Reverse transcription Real Time quantitative PCR (RT-qPCR)

Conjunctival models were lysed in 350 µl TRK lysis buffer (VWR International, Radnor, Pennsylvania, USA) supplemented with 20 µl/mL b-Mercaptoethanol and homogenized in a Tissue lyser (Qiagen, Hilden, Germany). Total RNA was isolated using the peqGold Total RNA Kit (VWR International, Radnor, Pennsylvania, USA). 500 ng of RNA was transcribed into cDNA using the iScript™ cDNA Synthesis Kit (Bio-Rad, Hercules, California, USA). RT-qPCR was performed using the SsoAdvanced Universal SYBR Green Supermix (Bio-Rad, Hercules, California, USA) and the QuantStudio™ 7 Flex system (Thermo Fisher, Waltham, USA). All runs were performed with 60 °C annealing temperature. Gene expression was calculated using the ΔΔCT-method with *18s* as housekeeper reference. Primer sequences (Eurofins Scientific, Luxembourg) were used as stated in Table 1.

**Table 1.**
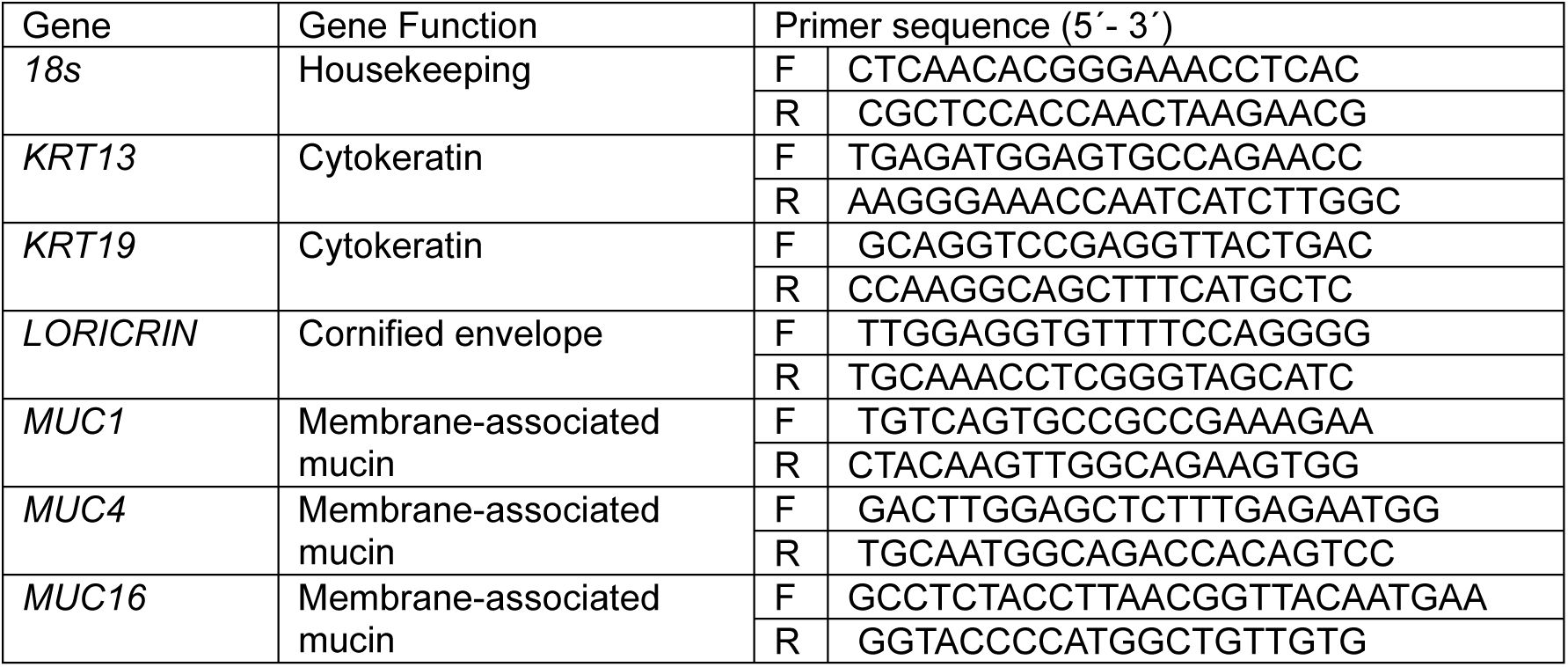
Primers used for RT-qPCR.

### ELISA

The concentration of secreted MUC5AC was assessed via ELISA of cell culture supernatant samples from day 15 of model culture with fresh culture medium as a control. Samples were stored at -80°C until further use. The assays were performed according to the manufacturer’s instructions (Human MUC5AC ELISA Kit (colorimetric), bio-techne, Minneapolis, MN, USA). Colorimetric measurements were performed on a TECAN Spark system (Tecan Group, Männedorf, Switzerland). MUC5AC concentrations were determined from a standard curve.

### Statistical analysis

Graphpad Prism 10 software (Graphpad Software Inc., La Jolla, CA, USA) was used for statistical analyses. Data sets were tested for normality via Shapiro-Wilk tests. Quantitative OCT analysis was evaluated via Two-way ANOVA. For RT-qPCR analyses, normally distributed data was analyzed via parametric One-way ANOVA with Tukey’s multiple comparisons test, while non-normally distributed data sets were analyzed via Kruskal-Wallis tests with Dunn’s multiple comparisons test. Values of *p* < 0.05 were considered significant.

## ACKNOWLEDGMENTS

The authors thank the Department of Ophthalmology of the University Hospital Würzburg for kindly providing biological material used in this study. The authors further thank Barbara Bayer, Annika Baumann, and Alevtina Höchner for their excellent technical support. This study was funded by the European Union within the frame of the Horizon Europe project: HORIZON-HLTH-2023-TOOL-05-01 under grant agreement No.101137315.

